# A high performing brain-machine interface driven by low-frequency local field potentials alone and together with spikes

**DOI:** 10.1101/015750

**Authors:** Sergey D. Stavisky, Jonathan C. Kao, Paul Nuyujukian, Stephen I. Ryu, Krishna V. Shenoy

## Abstract

**Objective:** Brain-machine interfaces (BMIs) seek to enable people with movement disabilities to directly control prosthetic systems with their neural activity. Current high performance BMIs are driven by action potentials (spikes), but access to this signal often diminishes as sensors degrade over time. Decoding local field potentials (LFPs) as an alternative or complementary BMI control signal may improve performance when there is a paucity of spike signals. To date only a small handful of LFP decoding methods have been tested online; there remains a need to test different LFP decoding approaches and improve LFP-driven performance. There has also not been a reported demonstration of a hybrid BMI that decodes kinematics from both LFP and spikes. Here we first evaluate a BMI driven by the local motor potential (LMP), a low-pass filtered time-domain LFP amplitude feature. We then combine decoding of both LMP and spikes to implement a hybrid BMI.

**Approach:** Spikes and LFP were recorded from two macaques implanted with multielectrode arrays in primary and premotor cortex while they performed a reaching task. We then evaluated closed-loop BMI control using biomimetic decoders driven by LMP, spikes, or both signals together.

**Main Results:** LMP decoding enabled quick and accurate cursor control which surpassed previously reported LFP BMI performance. Hybrid decoding of both spikes and LMP improved performance when spikes signal quality was mediocre to poor.

**Significance:** These findings show that LMP is an effective BMI control signal which requires minimal power to extract and can substitute for or augment impoverished spikes signals. Use of this signal may lengthen the useful lifespan of BMIs and is therefore an important step towards clinically viable BMIs.

**Suggested PACS:**

87.19.L-Neuroscience
87.19.lu motor systems
87.19.rs Movement
87.19.R-Mechanical and electrical properties of tissues and organs
87.85.E-Neural Prosthetics
87.85.Wc Neural engineering
87.85.dd brain-machine interface
87.85.Ng Biological signal processing

**IOP Subjects:**

Medical Physics, Biological Physics

## Introduction

Motor brain-machine interfaces (BMIs) seek to restore movement to individuals with movement disorders by decoding movement intention from the brain in order to directly control a prosthetic device. To date, the highest-performing BMIs have been driven by spike activity recorded with intracortical multielectrode arrays; these have enabled non-human primates to accurately control computer cursors [1–3], robotic limbs [4,5], or the subject’s own musculature [6]. This research is now being translated to early-stage clinical studies in people with paralysis [7–9]. A critical challenge that must be overcome to enable clinically viable BMIs is to improve the device’s useful lifespan by maintaining high performance over a long period of time, thereby improving its risk-benefit balance [10]. Chronically implanted sensors often degrade over time and gradually lose their ability to record action potentials [11–14]. One approach to mitigate this is to decode multiunit spikes instead of well-isolated single unit activity [3,8,13,15,16]. An alternative – or complementary – strategy is to make use of neural signals other than spikes that contain information about movement intention and are available from the same sensors. The local field potential (LFP) is such a signal: it is obtained by low-pass filtering the same raw voltage signal from which spikes are high-pass filtered, and it carries information about the kinematics of planned [17] and executed [17–21] reaching movements.

Despite many offline LFP decoding studies [17–30] and a few closed-loop demonstrations [31,32], only very recently have there been reports of effective closed-loop cursor control driven by LFP [33,34]. While encouraging, the performance achieved by these first forays into LFP-driven BMIs is low compared to spike-driven performance and leaves open the question of how viable LFP is as an alternative and complimentary control signal. This study aims to address this gap by showing considerably improved LFP-driven performance and by showing, for the first time, that under certain conditions LFP can be beneficially combined with available spikes to improve closed-loop BMI control. LFP can be processed into a variety of different features, and these two previous studies made specific design choices to decode LFP power in multiple frequency bands as well as – in [33] – low-frequency LFP amplitude known as local motor potential (LMP, which has also been referred to as movement-evoked potential in earlier studies). Closed-loop performance using other choices of LFP features has yet to be characterized. Based on the results of our own offline evaluation of decoder performance using various LFP features, we identified that the LMP was the best candidate feature. We subsequently evaluated the closed-loop performance of a BMI design that differs from that of previous studies by only extracting from each channel a half-wave rectified variant of the LMP, rather than the unrectified LMP or LFP power in various frequency bands. Two macaques successfully used this LMP-driven BMI to perform a 2D target acquisition task and, to our knowledge, demonstrated higher performance than in any previously reported online LFP BMI study.

After showing that this LMP feature is an effective alternative BMI control signal, we set out to test whether combining LMP and spikes decoding could improve closed-loop performance. LFP reflects a spatial averaging of synaptic and other currents in the vicinity of the electrode and carries information that is distinct from spikes [35–38]. Furthermore, LFP can be informative even on electrodes which do not record spikes [22,23,29,39,40]. For these reasons it has long been hoped that decoding LFPs could augment BMIs, especially in cases where few channels have spikes available. We report here the first use of a hybrid LFP and spikes decoder for continuous BMI cursor control and show that this approach can indeed improve performance compared to spikes-only decoding. The benefit of hybrid decoding increased as we reduced the number of channels with informative spike activity. However, we also found that in our second monkey, who had more electrodes recording tuned spikes activity and worse overall LMP-driven performance, hybrid decoding was only beneficial if spikes were made unavailable from the majority of electrodes.

A preliminary report of this study appeared in a conference proceeding [41].

## Methods

### Subjects and neural recording

All procedures and experiments were approved by the Stanford University Institutional Animal Care and Use Committee. Experiments were conducted with adult male rhesus macaques (monkeys ‘R’ and ‘J’) previously implanted with two 96-channel multielectrode arrays (1 mm electrodes spaced 400 μm apart; Blackrock Microsystems, Salt Lake City, UT) using standard neurosurgical techniques as described in [42]. The arrays were placed in the primary motor cortex (M1) and dorsal premotor cortex (PMd) contralateral to the reaching arm. Monkey R’s arrays were implanted 24-25 months prior to these closed-loop and ‘sedated reaching’ (see next paragraph) experiments and 16-20 months prior to collection of the offline decode datasets. Monkey J’s arrays were implanted 49-50 months before the closed-loop and sedated reaching experiments and 24 months prior to the offline datasets.

We used an “arm not restrained, not visible” animal model for BMI study, meaning that the monkey’s reaching arm was unrestrained, but visually occluded by the display, during both arm controlled and BMI controlled tasks [43]. We believe that this is a good BMI animal model because it allows training biomimetic decoders from natural reach data, and subsequently allows the animal to control the BMI cursor by generating whatever neural activity he wants without an additional constraint that this activity must not cause movement of the arm [44]. As a control experiment to test if LMP could be merely an arm movement artefact, we also collected datasets in which each monkey was sedated while an experimenter performed the Radial 8 Task by moving the monkey’s hand from target to sequential target. The animal was sedated with ketamine 2 mg/kg plus dexmedetomidine 0.04 mg/kg IM and positioned in the experimental rig in the same fashion as during awake-behaving experimental sessions. The experimenter was electrically insulated from the animal and moved the monkey’s hand so as to approximately match the animal’s typical reach trajectories; meanwhile, the behavioral task software was running, thereby allowing for identical analysis of this control experiment data. Neural and kinematics recording, signal processing, and subsequent data analysis were identical to that of our standard awake-behaving reaching experiments.

### Behavioral tasks

The monkeys were trained to sit head-restrained in a primate chair and perform reaching tasks with one hand, as shown in figure 1a. A virtual cursor and target were displayed in a 3D environment (MSMS, MDDF, USC, CA, USA) in front of the monkey by two 120 Hz LCD displays visually fused by mirrors. However, all of the cursor tasks in this study were confined to a 2D vertical plane. Custom experiment control software written on an xPC Target platform (Simulink, The Mathworks, Natick, MA, USA) controlled the behavioral task and updated the monitors with latency of 7 ± 4 ms. The monkey controlled the cursor position either by moving his hand, which was tracked with an infrared reflective bead-tracking system at 60 Hz (Polaris, Northern Digital, Canada), or with a velocity command from decoded neural activity (see below). In all tasks, the monkey earned a liquid reward after each successful target acquisition.

At the beginning of each experimental session the monkey performed a block of the Radial 8 Task. In this task, the location of the target alternated on each trial between the center of the workspace and a peripheral location pseudorandomly chosen from eight locations equally spaced along a circle of radius 12 cm. To successfully complete a trial, the monkey had to keep the cursor inside a 4 × 4 cm acquisition area around the target for a ‘target hold period’ of 500 consecutive ms. A new center trial immediately followed a successful trial or after failure to acquire the target within a 2 s time limit. These data were used for decoder training during closed-loop experiments, and previously collected Radial 8 Task datasets were used for offline analyses comparing different LFP-derived neural features.

Unless stated otherwise, we evaluated closed-loop performance on the Continuous Random Target Task. At the start of each trial, the target appeared in a random location anywhere in a 20 × 20 cm box centered in a bounded workspace that was itself 40 × 32 cm. The monkey had up to 8 s to acquire the target by keeping the cursor within a 5 × 5 cm acquisition area for a target hold period of 500 ms (monkey R) or 300 ms (monkey J).

We also evaluated performance on two additional tasks to compare our BMI to those of prior LFP studies. The first was a variant of the Continuous Random Target Task with target size reduced to 4 × 4 cm, trial time limit of 10 s, and hold time reduced to 100 ms, as in [33]. Due to differences in animal training, unlike in that study we did not fail a trial if the cursor left the target within 100 ms and later came back to make a successful acquisition. Instead, we post-hoc labeled these trials as failures. We note that this is a more punitive metric of our subjects’ performance because the animals did not have feedback to strongly incentivize them to avoid leaving the target. The second task was a variant of the Radial 8 Task similar to [34] with 3.4 cm wide targets located around a 13 cm diameter circle. Acquiring the center target initiated a trial but only reaches to the peripheral targets were analyzed. We based our variant of the So and colleagues study’s task on an earlier report [45] and therefore used a 300 ms hold time and 8 second time limit. We note that in [34] the hold time was actually 400 ms and the time limit was 10 seconds.

### Signal processing

Voltage from each of 192 electrodes (henceforth referred to as channels) was initially sampled at 30 ksps by a Cerebus system (Blackrock Microsystems) and analog filtered from 0.3 to 7500 Hz with a 3^rd^ order Butterworth filter (figure 1b). To extract multiunit spikes (also known as “threshold crossings”), the raw neural data from each channel was digitally filtered from 250 to 7500 Hz (3^rd^ order Butterworth). A spike was detected whenever the signal went below a threshold set at the beginning of each recording session to be -4.5 times the root mean squared value of the channel’s voltage [13]. Spike counts in non-overlapping bins from each channel formed a vector of spike features computed at each decoder time step. For all offline analyses this decoder time step was 50 ms. Prior studies have found that such short time steps lead to better closed-loop control than the longer values more commonly used in offline decoding studies [3,46]. For monkey R’s closed-loop experiments we used the same 50 ms time step, while for monkey J we used a time step of 25 ms based on the observation that his spikes-only performance improved with this even shorter time step. Spikes extraction from disconnected electrodes or electrodes with no spikes was disabled; depending on the dataset, 168-174 channels recorded spikes in monkey R and 188-192 in monkey J.

LFP was obtained by digitally low-pass filtering the same raw neural data below 500 Hz (4^th^ order Butterworth) and then downsampling to 2 ksps. For our offline decoding analysis, we evaluated a number of candidate LFP frequency bands in addition to the time-domain local motor potential (LMP), which is described in the next paragraph. We created a vector of LFP power features at each 50 ms time step for each channel by taking the mean power of the causally bandpass filtered signal (3^rd^ order Butterworth) from the previous 50 ms of LFP. Because a 50 ms window does not capture a full period of the signal for bands with minimum frequency *f*_min_ below 20 Hz, for these features we used overlapping bins of 1/*f*_min_ seconds slid every 50 ms. The amplitude response of the filter cascades used to generate each candidate LFP feature, including the effects of the shared initial signal processing described above, are shown in figure 2b. These were computed by simulating the analog and digital filters applied by the Cerebus signal processor, and then analyzing the series of these filters, plus the LFP frequency band filter of interest, using MATLAB’s cascade and filter visualization (fvtool) tools.

To obtain the LMP feature from a given channel, LFP samples were first clipped at ±300 μV to mitigate intermittent noise bursts. The LMP was then computed as the sliding mean voltage from the past 50 ms of LFP (figure 1c). The LMP is thus the low-pass filtered LFP amplitude (figure 2b). For offline decoding the LMP was decoded directly as described above, but in pilot closed-loop studies in monkey R we found that an additional step was needed to enable effective continuous cursor control. We observed that on most channels, the negative voltage deflection at the start of the reach was followed by a positive ‘after-potential‘, as seen in the top panel of figure 1d. During closed-loop BMI use this led to the cursor ‘springing back’ in the opposite direction shortly after a movement was initiated. This markedly impaired performance (supplementary figure 2). We therefore applied a half-wave rectification step where all LMP input values below 0 μV were rectified while positive input values were set to 0, as illustrated in the bottom panel of figure 1d. While zeroing positive voltages on all channels worked well across the four arrays used in this study, we anticipate that in some situations the sign of this rectification step may need to be set on a channel-by-channel basis depending on each channel’s initial LMP polarity. To provide the reader with further intuition about the LMP signal, figure 2c shows the spectral content of the LMP used to control the BMI on two different electrodes from representative experimental sessions. These spectra were generated by applying the MATLAB *fft* function to the full duration of each electrode’s LMP during BMI task. The spectra were binned into 1 Hz bins, with figure tick locations correspond to the maximum frequency of each bin. The signal magnitude within each frequency bin was then normalized by total magnitude over all frequencies.

### Neural decoding

A velocity Kalman filter (KF) was used to decode cursor velocity from neural activity for both offline decoding and closed-loop BMI experiments. The details of the velocity KF as applied to BMIs are well described in [3,47,48]; here we enumerate specific design choices used for the present study. Briefly, the KF estimates the state of a linear Gaussian dynamical model where the kinematics state at time t, **x**(t) evolves from the previous time step’s state and relates to neural activity **y**(t) according to,

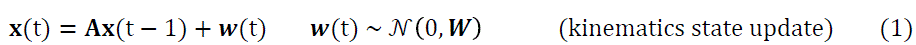

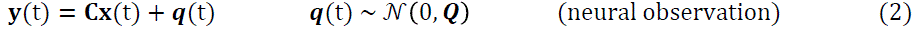

where **x**(t) is the velocity vector [v_x_(t) V_y_(t) 1]^T^ (the 1 is an offset term to accommodate non-zero-mean neural data), **y** (t) is a vector consisting of the neural feature(s) from every channel at time t, **A** is the matrix describing the linear kinematics dynamics, **C** is the observation matrix mapping kinematics state to neural activity, and **w**(t) and **q**(t) are Gaussian noise. **A**, **C**, **W**, and **Q** are fit from binned velocity and neural feature(s), from the start to the end of each successful trial in the training dataset. Once the model is learned, decoded kinematics can be inferred at every time step using the KF, which is a closed-form recursion. When regressing neural data against kinematics to fit **C**, we allowed the decoder to try different causal lags (i.e., neural data either aligned with or leading kinematics) for each type of neural feature: spikes, LMP, or LFP power in different frequency bands (which were computed using either 50 ms bins or, for lower frequency bands, longer bins slid every 50 ms). This lag was applied globally to all channels. For the final decoder we fit parameters with whichever lag minimized the residual. For LMP decoding the best lag was 100-125 ms (depending on the dataset), while for spikes decoding it was 50 ms in monkey R and 150 ms in monkey J. However, when fitting spikes decoders for closed-loop evaluation we used a 0 ms lag so as to be consistent with the methods used in [3,49]. We note that the offline spikes decoding difference between using the best lag and 0 lag was small and not significant: correlation between true and reconstructed velocity r = 0.73 vs. r = 0.72 for monkey R, and r = 0.80 vs. r = 0.77 for monkey J (p > 0.05 for a difference in means, paired *t*-test). For closed-loop LMP decoding, we only applied the 100-125 ms lag to the training data used to fit the models; no lag was used during closed-loop use. This choice follows with the general control systems design principle that minimizing control loop lag improves performance when feedback is used to change the control signal [50], as is the case for closed-loop BMI use. Reducing lag has been empirically shown to be best practice in the context of motor control [51] and BMI use specifically [3,46,52]. By applying a lag in the training data only, we were trying to better align neural correlates of movement intention with kinematics. During closed-loop use, we wanted the observation of similar neural activity to then generate intended cursor velocity with as little delay as possible.

We also applied an additional operation during LMP-driven BMI use to help stabilize the cursor in the workspace. A ‘centering velocity’ pointing towards the center of the workspace was always added to the decoded velocity, with magnitude 0.15 · *r*/s, where r is the distance from the cursor to the workspace center. Thus, the intervention was minimal when the cursor was near the workspace center but increased when the cursor was near the edges. We emphasize that this assistance is not dependent on target location; the centering velocity always moved the cursor towards coordinate (0,0), whereas targets could appear throughout the workspace. This was helpful because a velocity-only decoder can accrue a position bias over time; the centering velocity ameliorates this bias. The average centering velocity applied during LMP-driven cursor control was up and to the left in both monkeys (monkey R: -1.34 ± 0.19 mm/s x-velocity, 2.33 ± 0.30 mm/s y-velocity, grand mean ± std across datasets; monkey J: -1.14 ± 0.23 mm/s x-velocity, 3.32 ± 0.50 mm/s y-velocity. All y-velocity means and monkey R’s x-velocity mean were significantly different [p < 0.1] from zero by two-sided i-test).

This indicates that there was a down and to the right decoder bias. The centering velocity gain value of 0.15 was chosen based on a cursory parameter sweep during pilot experiments but was not thoroughly optimized, either in magnitude or in directionality (i.e., the centering velocity was symmetrical around the center and thus naïve to the particular subject’s typical bias); a more sophisticated bias correction scheme is a potential avenue for future work. We also sometimes applied this centering velocity when evaluating hybrid decoders in the channel-dropping experiment described later. To prevent this from potentially giving hybrid decoders an undue advantage over spikes-only decoders, we evaluated spikes-only decoders both with and without the same centering velocity and recorded the higher performance of the two. We note that this conservative methodology slightly favors the null hypothesis that hybrid decoding does not perform better than spikes-only decoding.

To evaluate whether decoding both LMP and spikes together would improve closed-loop performance, we chose to start with a state-of-the-art spikes decoder and then compare it to a hybrid spikes + LMP decoder. For the spikes-only decoder we used our previously reported Feedback Intention Trained Kalman filter (FIT-KF) [49]. The FIT-KF improves upon a standard velocity KF by adjusting kinematics of the training data to attempt to better match the subject’s true intent: cursor velocity is rotated to always point towards the target and is set to zero during target hold epochs. Unlike in [49], here we kept a number of design choices from the original ReFIT-KF decoder [3] from which FIT-KF is descendent: we trained the decoder using Radial 8 Task reaching data, and used data starting from the beginning of each trial.

The hybrid spikes + LMP decoder was built by combining the spikes FIT-KF and LMP velocity KF models, which were fit from the same training data. This decoder operated on a stacked feature vector y = [y_spikes_; y_LMP_] which was mapped to kinematics by a stacked C_hybrid_ = [C_spikes_; C_LMP_]. Since A_spikes_ was fit from intention-estimated velocities, it differed slightly from A_LMP_; we set A_hybrd_ = A_spikes_ and W_hybrid_ = W_spikes_. A combined covariance matrix Q_hybrid_ was then calculated from the training data and C_hybnd_, yielding the complete KF parameters {A_hybnd_, W_hybnd_, C_hybird_, Q_hybnd_}. Note that during closed-loop use, position feedback subtraction [3] was performed on the observed spike counts as in FIT-KF, but not on the observed LMP.

### Performance measures

The primary metrics we used to quantify closed-loop task performance on the Continuous Target Acquisition Task were success rate and normalized time to target (NTTT), which directly affect the animal’s rate of liquid reward. Success rate is defined as the number of trials in which the target was successfully acquired, divided by the total number of trials. NTTT is the time duration between trial start and successful target acquisition (not including the required target hold time), divided by the straight-line distance between this trial’s target center and the previous trial’s target center [53]. NTTT is only computed for consecutively successful trials. Unless otherwise stated, when comparing two different decoders, we first compared success rates using each decoder (binomial test). If success rates were not significantly different, we then compared the NTTT with two-tailed t-tests for a difference in means. A threshold of p < 0.01 was used for both comparisons. Chance performance levels were calculated under the null hypothesis that the cursor’s movements were not goal-directed towards acquiring the displayed target. We did this by replaying the time series of actual cursor trajectories recorded during the BMI session while randomly selecting virtual target locations drawn from the set of all targets presented during the experiment. A virtual trial ended successfully if the cursor stayed within the target for the requisite hold time, or unsuccessfully if the 8 s time limit was exceeded. At the end of each virtual trial the next random target was selected and the kinematics playback continued. Chance success rate and median NTTT were then calculated by averaging across 1,000 repeats of this target-shuffled simulation. This method of calculating chance performance differs from the two methods used in [33] and [34]; in particular, it yields a comparatively higher chance success rate when trial time limits are long, which is tempered by long chance times to target. We chose to use this method because it is less prone to underestimating chance success rates of undirected kinematics which nonetheless traverse much of the workspace.

We also computed three additional metrics of successful trials’ kinematics. Index of performance (IP) is a throughput metric for human-computer interface devices motivated by Fitts’ law [54] and is specified in the ISO 9241 standard [55].

It was adapted to two-dimensional tasks [56] and adopted by the BMI field (e.g., [3,11]). IP is calculated for each trial and rewards faster target acquisition while taking into account differences in target size and distance:

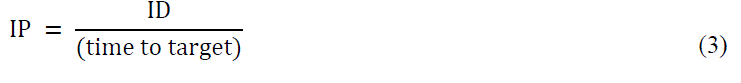

where ID is the index of difficulty which depends on this trial’s reach distance, *D* (i.e., Euclidean distance between the cursor’s position at the start of the trial and its position when the target is successfully held [11]), and the width of the target, *W*:

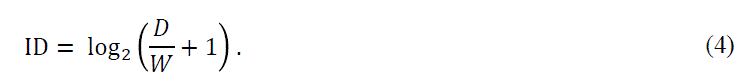

‘Dial-in time’ is calculated for each trial and measures the time between when the cursor first entered the target acquisition area and when it last entered the target area prior to successfully staying inside the target area for the requisite hold time [3]. A dial-in time of 0 ms means that the monkey reached and successfully held the cursor inside the target on his first attempt. A longer dial-in time signifies that the cursor left the target acquisition area too soon, which required the monkey to reacquire the target. Lower mean dial-in times signify better ability to stop the cursor over the target and hold it there.

Path length ratio (sometimes also referred to as ‘distance ratio’ or the inverse of ‘path efficiency’) is the ratio of the distance the cursor actually travelled from the beginning of the trial to when it entered the target acquisition area, divided by the straight-line distance between the cursor’s position at the start of the trial and the edge of the target acquisition area. Path length ratios closer to 1 signify straighter cursor trajectories.

### Channel preferred direction and decoder contribution

Data from the hand-controlled Radial 8 Task were used to compute the directional tuning of LMP, as well as each channel’s contribution to both the LMP and spikes velocity decoders fit from this data. To compute a simple measure of preferred direction and tuning significance for a given channel, we first averaged this channel’s LMP over an analysis epoch spanning 200 to 600 ms into each trial. For context, the median and standard deviation of movement start times (defined as when hand speed reached 10% of the trial’s peak speed) was 144 ± 90.3 ms for monkey R and 222 ± 68.6 ms for monkey J. The hand entered the target after 511 ± 196 ms (monkey R) and 495 ± 141 ms (monkey J). The channel’s LMP averaged over the analysis epoch yielded a single data point per trial. We then grouped trials by reach target and performed a one-way ANOVA to test whether LMP was significantly different for reaches towards at least one of the eight targets. Our measure of each channel’s preferred direction was computed by summing eight individual vectors that pointed in the prompted reach directions, where each vector’s length equaled this channel’s average LMP across all trials where the monkey reached in that direction.

The magnitude of each channel’s contribution to the decoder [16] was used both for determining channel dropping order and for comparing the cumulative channel contribution curves of spikes and LMP decoders. To find each channel’s contribution to a particular decoder we first converted that Kalman filter to a closed-form steady-state [57],

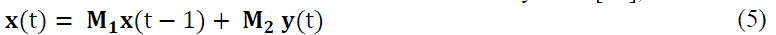

where the j’th column of **M**_2_ is the decoder weight vector **W**_j_ = [w_x,j_ W_y,j_ 0]T that describes how y_j_(t), the neural activity observed on channel j, contributes to x-and y-velocity. Channel j’s decoder contribution is then computed as that channel’s average effect on cursor velocity:

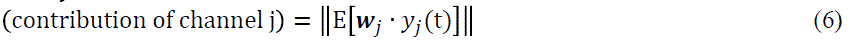

where the expectation is taken over time. A channel’s relative contribution to the decoder is then calculated by dividing that channel’s contribution by the sum of all channels’ contributions. In the channel dropping experiments, for each condition we removed the top *n* channels ranked in order of most to least contribution to the full-channel spikes decoder.

### Comparing the contribution of LMP and spikes in the hybrid decoder

We dissected LMP’s and spikes’ respective contributions in the hybrid decoder using the steady-state decoder form described above. At each 50 ms time step, the instantaneous contribution to decoded velocity (“neural push”) **x**(t) was divided into **X**_spikes_(t) = **M**_2_ · [y_spikes_(t); **0**] and **X**_LMP_(t) = **M**_2_ · [**0**; **y**_LMP_(t)], where **y**_spikes_ (t) and **y**_LMP_ (t) are the spikes and LMP feature vectors, respectively, and **y**_spikes_(t) incorporates the position subtraction operation of the ReFIT-KF [3]. The relative contribution of each signal (LMP or spikes) was then computed by dividing the magnitude of its velocity contribution by the sum of both signals’ velocity contribution magnitudes. This metric was averaged across all time steps from 200 ms after target presentation until when the cursor first acquired the target.

We also used the partial decoded velocities **x**_spikes_(t) and **x**_LMP_(t) to find the angular error between the direction pointing directly from the cursor towards the target and the direction that each signal pushed the cursor. We report the mean angular error for LMP and spikes averaged across all time steps included in the decoded velocity contribution analysis described above. This decode error metric assumes that at each time step the animal’s true intended velocity points directly from the current cursor position towards the target. However, due to delays in the control loop (of both biological and technical origin), the neural signals observed at time *t* likely reflect an intended velocity towards the target from the cursor’s earlier position at time *t* – τ. We therefore also computed angular error as described above across different lag values of τ. We were particularly interested in the difference in error-minimizing τ between the spikes and LMP signals because this could reveal differences in how quickly the animal modulates each type of control signal. For this analysis we used each trial’s data starting from 300 ms after target presentation (rather than 200 ms, as above) to allow testing a wider range of τ.

For the analyses of closed-loop hybrid decoder use, all experimental sessions evaluating the full-channel hybrid decoder were included. For the matched offline hybrid decoding comparisons of figure 7(e,f), we analyzed data from experimental sessions where the monkey performed the same Continuous Random Target Task with his hand while neural data was recorded (812 trials from 2 sessions for monkey R, 1721 trials from 4 sessions for monkey J).

## Results

### Selection of local motor potential as a control signal based on offline evaluation of candidate LFP features

In order to choose which LFP-derived feature(s) to evaluate as the control signal for closed-loop BMI experiments, we first tested a number of neural features offline to determine how well the velocity of point to point arm reaches could be decoded from each feature. We compared spike counts in non-overlapping 50 ms bins, the local motor potential (LMP, i.e., low-pass filtered LFP amplitude), and LFP power in the 1-4 Hz, 3-10 Hz, 12-23 Hz, 27-38 Hz, and 50-300 Hz bands. We also tested decode performance using combinations of spike counts and one or more LFP-derived features. For this analysis we used five datasets of the Radial 8 Task from each of two adult male rhesus macaques. A velocity Kalman filter was trained to decode 2D hand endpoint velocity from each type of neural feature or combination of features. Neural data came from 192-channel multielectrode recordings from contralateral M1 and PMd. Half of each dataset’s ∼500 trials were used for training the decoder and the remaining half were used to test how accurately the decoder reconstructed the true hand velocity. Kinematics and neural features were binned every 50 ms, and decode performance using different lags (i.e., neural data aligned to or leading kinematics) was evaluated; we report how accurately the testing data was decoded using the best lag for each particular feature as fit from the training data.

Figure 2 shows the average of the Pearson’s linear correlation coefficients between true and reconstructed x-and y-velocities using each neural feature(s). We found that LMP outperformed the various LFP power features, and was the only LFP-derived feature to match or exceed the offline decoding accuracy of spikes (monkey R spikes r = 0.73 ± 0.022 [mean ± standard deviation], LMP r = 0.78 ± 0.010, p = 0.007 [paired t-test for a difference in means]; monkey J spikes r = 0.80 ± 0.032, LMP r = 0.77 ± 0.005, p = 0.065). Combining LMP with spikes improved offline decoding accuracy in both animals compared to decoding spikes alone (monkey R Δr = 0.11, p < 0.001; monkey J Δr = 0.054, p = 0.003). Although decoding both gamma LFP power (50-300 Hz) and spikes together also led to a small increase in accuracy, this effect was smaller than when combing LMP and spikes. Furthermore, adding both LMP and gamma to spikes did not substantially improve the decode accuracy over just using spikes + LMP (monkey R Δr = 0.0026, p = 0.034; monkey J Δr = 0.0002, p = 0.78). LMP was also a better offline decode feature than signal amplitude in the other bands, or acausally filtered power in the other bands (supplementary figure 1).

**Figure 1.**
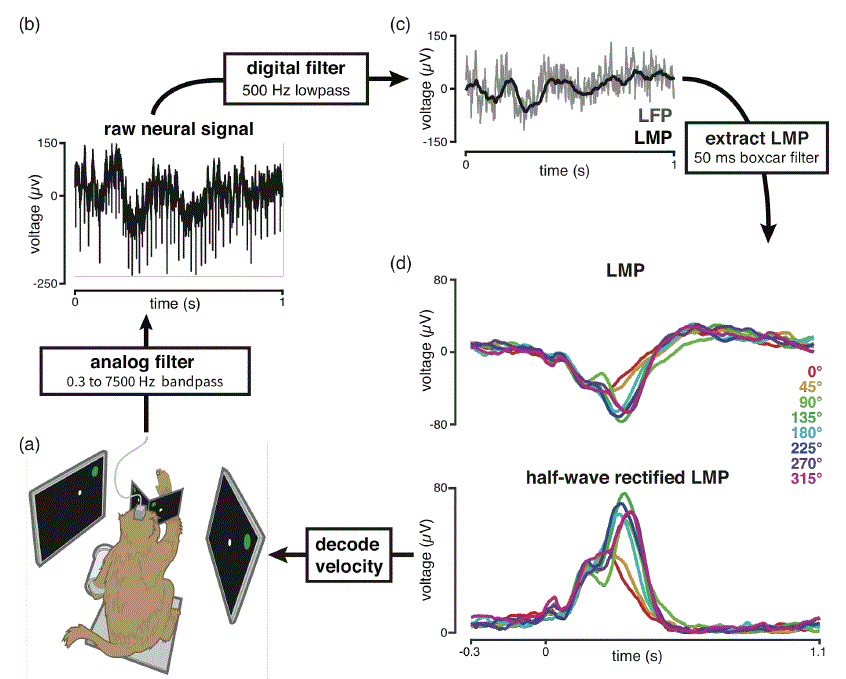
Overview of the signal processing of the local motor potential (LMP) driven BMI system. (a) A macaque sat in a primate chair and observed a cursor and target in a virtual-reality environment. During arm control, he controlled the cursor position by reaching with his unrestrained arm. During BMI control, the cursor was driven by a velocity command decoded from neural activity. (b) Neural signals were recorded from 96-channel multielectrode arrays in M1 and PMd. Electrical signals were amplified, analog filtered from 0.3 to 7500 Hz, and digitized. A 1 s sample of this raw signal is shown; both lower frequency components and high frequency multiunit spikes are visible. (c) The LFP (shown in grey) was then obtained by low-pass filtering the raw digital signal below 500 Hz. LMP, shown in black, was extracted by boxcar filtering the most recent 50 ms of LFP. (d) (Top) Example LMPs trial-averaged over 32 arm reaches to each of 8 peripheral targets. Trace color corresponds to reaches to targets in the specified direction. The target was presented at time = 0. (Bottom) Before decoding, LMP was half-wave rectified: negative voltage samples were rectified while positive voltage samples were set to 0. Dataset R.2013.07.30, channel 1 (M1 array).

**Figure 2.**
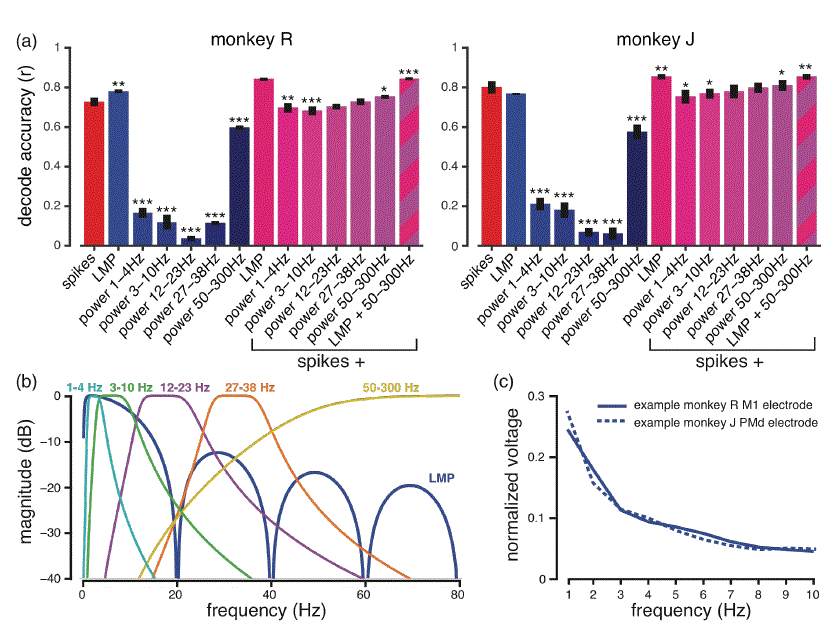
Comparison of offline decode accuracy using different neural signal features. (a) Two-dimensional hand velocity from Radial 8 Task reaches was reconstructed offline with a Kalman filter that decoded either multiunit threshold crossing spikes (red), various LFP features (blue), or the combination of spikes and one (violet) or two (striped) LFP features. The Pearson correlation (r) between true and decoded velocity was averaged across all trials and both x-and y-dimensions. We show the grand mean and standard error of measurement across five datasets for each monkey. Stars denote signals or combinations of signals whose decode performance differed significantly from that of decoding spikes only (paired *t*-test, * p < 0.05, ** p < 0.01, *** p < 0.001). (b) Amplitude response of the filter cascade used to generate each of the evaluated LFP frequency bands. The periodic dip in the response of LMP is due to the 50 ms boxcar filtering operation. The high frequency cutoff of the 50-300 Hz filter extends beyond the panel; it is flat until 200 Hz and rolls off to -4.7 dB at 300 Hz. (c) To illustrate the actual spectral content of the half-wave rectified LMP signal, we have plotted the frequency spectra of this feature during closed-loop use of the LMP decoder from a representative electrode from each monkey. Datasets R.2013.09.09 channel 1 and J. 2013.09.11 channel 122.

Velocity could be decoded from LMP because this signal evolved differently over time depending on which direction the animal was reaching. This can be seen in figure 1d, which shows larger amplitude LMP deflections during reaches towards a subset of the 8 targets on an example M1 electrode. Across electrodes and monkeys we observed a variety of LMP waveforms in terms of shape, number of peaks, and tuning for velocity (not shown). Figure 3 shows that the LMP recorded on most electrodes was significantly tuned for reach direction, and that the preferred directions (PDs) of LMP recorded on nearby electrodes tended to have similar tuning. Despite these local correlations, across all electrodes the PDs spanned both dimensions required for 2D cursor control. We note that monkey R’s M1 array is an example of an array that records very poor spikes signals (offline decode accuracy of r = 0.11 ± 0.042 [mean ± standard deviation] using just this array’s spikes to perform the same analysis as in Figure 2), but has significant LMP tuning on a number of electrodes. When decoding LMP from only this array, we observed an improved offline decode accuracy of r = 0.46 ± 0.029. Based on these offline results we decided to evaluate LMP alone and spikes + LMP together as control signals for a closed-loop BMI.

**Figure 3.**
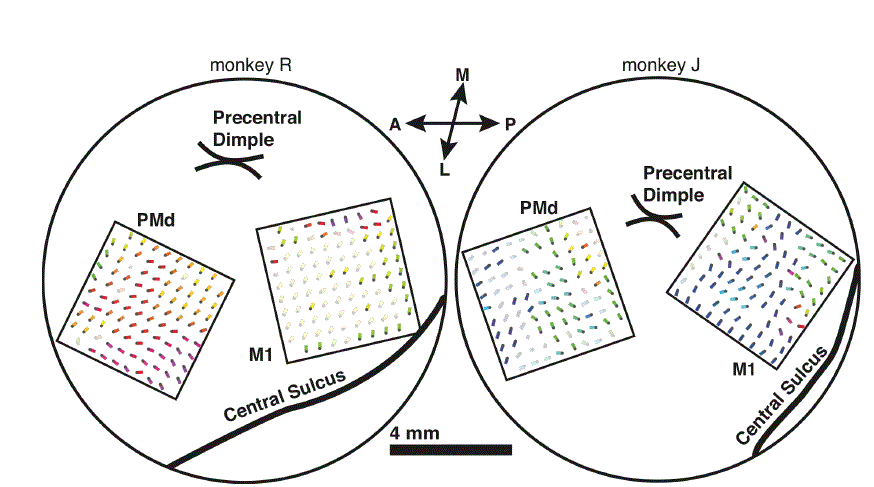
Directional tuning of LMP for each channel of the multielectrode arrays overlaid on electrode location in motor cortex. The locations of the two arrays in each monkey are drawn along with major anatomical landmarks based on intraoperative surgical photographs. Each dot with a coloured vector coming out of it corresponds to an electrode and the preferred direction (PD) of the LMP feature recorded on that electrode, as computed from 250 center out arm reaches. Both the vector’s colour and angle correspond to PD, while the vector length signifies 1- (*p* value! of tuning (ANOVA). Channels that were not significantly tuned (p > 0.01) are shown in greyed out colour. Note that the PDs shown are in the coordinate system of the two-dimensional task and are unrelated to the anatomical coordinate system used to map the physical location of each electrode in the brain. PMd = dorsal premotor cortex, Ml = primary motor cortex, A = anterior, P = posterior, L = lateral, M = medial. Monkey R’s Ml array quality was poor both for LMP and especially for spikes even from the earliest post-implant recordings. Datasets R.2013.09.09 & J.2013.09.11.

### Control experiment showing that a sedated animal’s passively moved arm kinematics cannot be decoded from LMP

Prior to conducting closed-loop experiments, we first wanted to convince ourselves that the LMP we were decoding was not artefactually correlated with the monkey’s arm movement due to factors such as microphonic pickup of electronic noise in the room (despite heavily electromagnetically shielded walls) or mechanical vibration of the recording equipment (despite the ruggedly built experimental apparatus being connected to massive load-bearing metal walls via structural metal). Both of these noise sources could appear to be ‘tuned’ to reach kinematics because arm movements can cause very small but specific movements of the experimental apparatus. We therefore conducted a control experiment in which similar arm movements were made without a corresponding neural movement intention. We sedated the monkey with ketamine and dexmedetomidine, gently placed him in the primate chair, and recorded neural signals while the monkey’s hand was moved by a human experimenter to complete the same Radial 8 Task. Mean hand velocity reconstruction accuracy using LMP dropped from r = 0.82 (monkey R), r = 0.79 (monkey J) when the animals were awake and making volitional reaches, to r = 0.05 (monkey R) and r = 0.01 (monkey J) using the sedated control data collected later the same day. This indicates that the LMP signal is not an artefact related to the monkey’s arm movements.

### Closed-loop BMI performance driven by LMP

Both monkeys demonstrated high performance using an LMP-driven BMI. For each experimental session, we trained a velocity Kalman filter using LMP from ∼500 arm reaches. We evaluated performance by measuring the success rate, normalized time to target, and path length ratio on a Continuous Random Target Task in which 5 × 5 cm targets appeared anywhere in a 20 × 20 cm workspace and had to be acquired within 8 seconds (figure 4). Monkey R did the task at a 99% success rate (11,429 trials across 6 experiment sessions) with 0.08 s/cm median normalized time to target and median path length ratio of 2.09. Supplementary Movie 1 shows a representative one minute of LMP decoder use by monkey R. Monkey J had an 86% success rate using the LMP decoder across 2,065 trials over 8 sessions, with median normalized time to target of 0.095 s/cm and path length ratio 2.47. Chance performance levels were 21% success rate, 0.33 s/cm time to target for monkey R, and 34% success rate, 0.31 s/cm time to target for monkey J. Note that we had given monkey J an easier task (300 ms target hold time compared to monkey R’s 500 ms hold time) to keep him engaged in the task because he had worse LMP-driven BMI performance.

Since there have only been a few published online LFP BMI studies, we were able to also measure performance on tasks closely matched to those used in two prior reports to allow for a more direct comparison. When tested on a Radial 8 Task similar to that of [34], monkeys R and J achieved considerably higher success rates: 98% for monkey R and 85% for monkey J (supplementary figure 3), compared to 78% and 72% success rates reported in [34]. In terms of successful target acquisitions per minute, monkey R’s 16.1 targets/min and monkey J’s 9.8 targets/min compare favorably to the 10.6 and 5.8 targets/min in [34]. We also had both monkeys perform a Continuous Random Target Task with smaller targets and 100 ms hold times as in [33] (supplementary figure 4). Monkey R performed this task with 79% grand mean success rate, normalized time to target = 0.070 s/cm, index of performance = 2.71 bits/s, and path length ratio = 1.53. Monkey J’s metrics were: 73% success rate, normalized time to target = 0.090 s/cm, index of performance = 2.16 bits/s, and path length ratio = 1.79. Compared to [33], this represents a slightly better success rate and over twice as fast target acquisition times. Note that these success rates are lower than in our standard Continuous Random Target Task because we posthoc counted as failures those trials in which the cursor exited the target and then successfully reacquired it (which were successful as far as the monkey was concerned). We did this to be consistent with the methods used by Flint and colleagues; otherwise, the success rates would have been 99% and 86% for monkeys R and J, respectively.

Since we evaluated biomimetic decoders trained using arm reaching neural activity, we did not restrain the monkey’s contralateral (with respect to the arrays) arm during BMI use. We observed that both monkeys continued to move their arms during LMP-driven cursor control. The monkeys were not required to keep the reflective bead taped to their fingers visible to the tracking system when using the BMI; consequently, we were only able to quantify the relationship between hand and cursor movement for 78% (monkey R) and 53% (monkey J) of LMP-driven time samples. During these epochs, the grand mean correlation between hand and LMP-driven cursor velocities was r = 0.736 (p < 0.001) for monkey R and r = 0.668 (p < 0.001) for monkey J (averaging across x-and y-velocity, and across all Continuous Random Target Task sessions).

**Figure 4.**
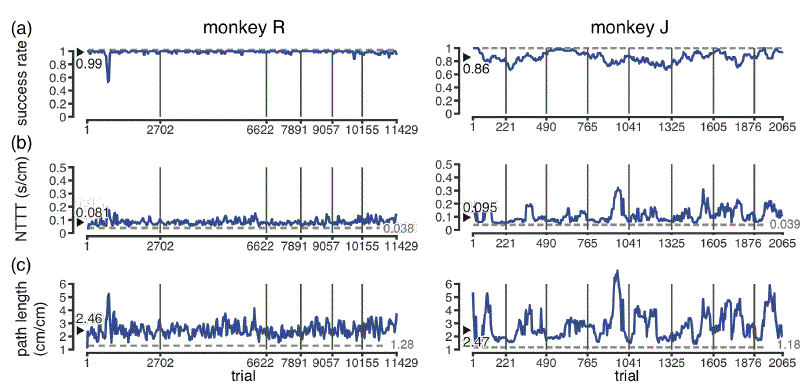
Performance using the LMP-driven BMI on the Continuous Random Target Task with 500 ms (monkey R) or 300 ms (monkey J) target hold times. Each performance metric is smoothed over the previous 50 trials and is shown across six and eight days’ experiments for monkeys R and J, respectively. Vertical gray lines separate trials from different days. Grand mean (success rate) or median (NTTT and path length) performance for each plot is marked with a black arrow and value. Dashed grey lines and grey numbers show performance on this task using hand control. (a) Success rate is the fraction of trials where the target was acquired within 8 seconds. (b) NorMalized time to target measures how long it took to successfully acquire the target, norMalized by the straight-line distance between the current and previous targets. The final hold time is not included in NTTT. (c) Path length ratio measures the distance the cursor travelled to acquire the target divided by the straight-line distance; closer to 1 is more direct (better).

### Fixed LMP decoders work across multiple days without retraining

We also evaluated how well a fixed LMP decoder performed without any retraining for just over three weeks. A decoder was trained on day 1 of these experiments and then loaded and evaluated for at least 200 trials at the beginning of subsequent days’ experimental sessions. No recalibration or re-normalization was done; the decoder weights and entire signal processing pipeline was identical between collection of the training dataset on day 1 and during evaluation of the fixed decoder. The fixed LMP decoder enabled both monkeys to perform the Continuous Random Target Task across the entire tested period (figure 5a-d), although with worse performance than using same day-trained decoders. Monkey R’s daily success rates using the day 1 decoder on subsequent days ranged from 90.4% to 99.8%, with an accuracy of 93.0% on the last day tested (day 22), while Monkey J’s daily success rate ranged from 69.8% to 86.4%, and was 72.7% on day

To investigate whether the ability to use a fixed decoder over several weeks was due to stable LMP signals or the animal adapting to a potentially mismatched decoder during closed-loop BMI use, we compared the fixed decoder’s closed-loop performance with its offline decode accuracy. We performed an offline decode of hand velocities recorded during Radial 8 Task arm reaches at the start of each of these experimental sessions using either the fixed day 1 decoder or a decoder trained from that same day’s reaches. The fixed day 1 decoder was only slightly worse than the same day decoder (figure 5e), suggesting that the LMP signals were largely stable across days. For monkey J, who exhibited a wider divergence in BMI performance between the fixed and same-day decoders, closed-loop day 1 decoder success rates correlated with the day 1 decoder’s offline accuracy (r = 0.79, p < 0.05, Pearson correlation). For monkey R, who had little variation in fixed decoder offline accuracy, this regression was not significant against any of the closed-loop performance metrics.

**Figure 5.**
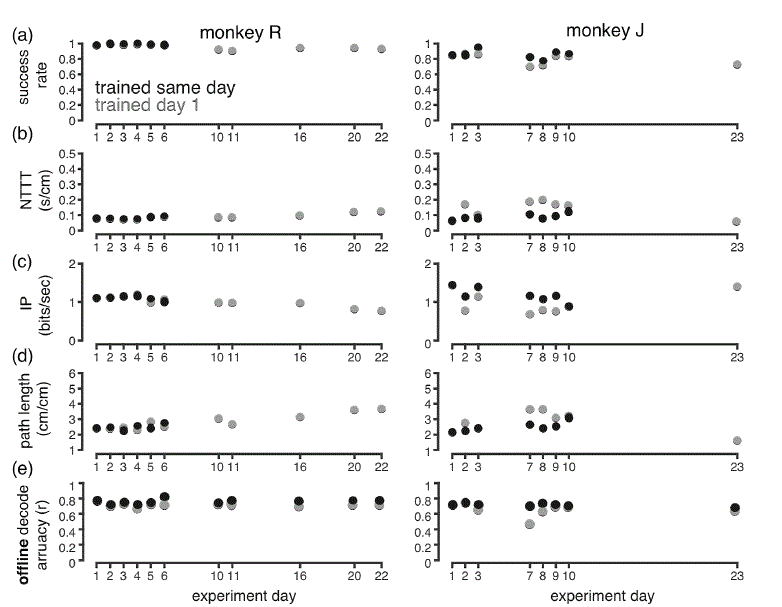
(a-d) Closed-loop performance of a fixed LMP decoder trained on day 1 of the experiment and then evaluated on subsequent days. Each grey point shows performance of this fixed decoder for each subsequent day’s experiment. On some days we also trained a new decoder from arm reaches; the performance of this same-day trained decoder is shown in black. On days 2-6 for monkey R the same day and day 1 decoders performed similarly, such that the black points obscure the grey points. Same day performance corresponds to the data shown in Figure 4. (a) Mean success rate. (b) Median norMalized time to target. (c) Median Index of Performance (IP), also known as Fitts bits per second. Faster acquisition times and/or accomplishing a harder task yields higher IP. (d) Median path length ratio. (e) Offline decode accuracy when predicting each day’s arm reaching hand velocities from LMP using either the fixed day 1 decoder (grey) or a decoder trained from that day’s data with 10-fold cross-validation (black).

A potential concern is that the monkey’s familiarity with the fixed decoder could have interfered with his ability to control new decoders provided on subsequent days. Aside from a brief pause of the task, no explicit cue was provided to indicate that the decoder had been switched. Nonetheless, we found that the monkeys rapidly adjusted to each day’s newly trained decoder after switching from the fixed decoder. The data shown in figure 4 begin following a brief (10-20 trial) ‘transition’ period after the switch from the fixed decoder, and do not show within-session improvement that would indicate overcoming interference from the first decoder.

### BMI control using a hybrid spikes and LMP decoder

We next evaluated whether simultaneously decoding both LMP and spikes would improve performance compared to a BMI driven by spikes alone. Figure 6 shows aggregated data from eight datasets in monkey R and five datasets in monkey J where the hybrid decoder and spikes-only decoders were evaluated in an interleaved ‘ABAB’ block format. When LMP features were added to spikes, the hybrid decoder outperformed the spikes-only decoder in monkey R (12% improvement in mean normalized time to target averaged over all datasets, p < 0.001; difference significant at p < 0.01 in 5/8 individual sessions). In Monkey J, however, the hybrid decoder was slightly worse than the spikes-only decoder (8% slower normalized times to target, p = 0.007, difference not significant at p < 0.01 in any individual session).

**Figure 6.**
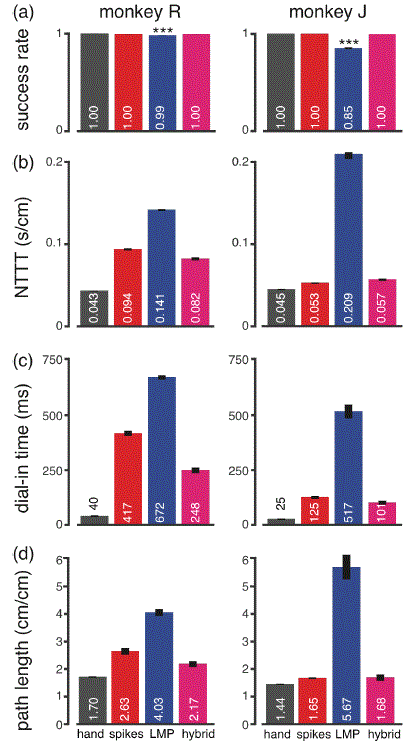
Comparison of spikes-only, LMP-only, and hybrid spikes + LMP decoders. Mean and SEM of each performance metric was computed across trials from multiple datasets in which the Continuous Random Target Task was performed with either the hand (grey), or a BMI driven by spikes (red), LMP (blue), or hybrid spikes + LMP (violet). (a) Success rate. The LMP decoder was significantly worse than other control modes (p < 0.001). (b) NorMalized time to target. All differences significant at p < 0.01. (c) Dial in time. All differences significant at p < 0.05. (d) Path length ratio. All differences significant at p < 0.05 except monkey J spikes vs. Hybrid.

To provide context for these results, we also compared each monkey’s spikes-only and hybrid decoding performance to his performance on the same task using either his hand (3 and 4 datasets from monkey R and J, respectively) or the LMP-only decoder (9 and 7 datasets from monkey R and J, respectively); these data came from different experimental sessions. Figure 6 shows that in both monkeys, BMI performance was better using a spikes-only decoder than an LMP-only decoder. This difference was much larger for monkey J (298% faster normalized times to target) than monkey R (51% faster). Using an LMP-only decoder, both monkeys were slower to initially reach the target and were worse at holding the cursor inside the target acquisition area for the requisite hold time. Monkey R’s mean time to initial target entry was 50% slower for LMP than spikes (937 ms vs. 624 ms) and his dial-in time was 61% slower (LMP: 672 ms, spikes: 417 ms). Monkey J’s initial target entry was 273% slower for LMP than spikes (1789 ms vs. 479 ms), and his dial-in was 313% slower (LMP: 517 ms, spikes: 125 ms). All of these differences were significant at p < 0.001 (two-sided *t*-test). We also note that monkey J had very good spike-driven performance, acquiring targets only 18% slower than with his hand.

To better understand why closed-loop hybrid decoding only modestly helped in monkey R and did not help in monkey J, we compared how spikes and LMP each contributed to the hybrid decoder both during online BMI control and during offline decoding of neural data recorded while the monkey did the same Continuous Random Target Task with his hand. At each decode update time step, spike counts and LMP features from all channels were input to the decoder to generate the instantaneous velocity component (‘neural push‘) due to each signal (figure 7a). In both monkeys, LMP contributed more than spikes to the BMI cursor’s movement, accounting for 67% of the decoded velocity in monkey R and 54% in monkey J (figure 7b). This ruled out the possibility that LMP was simply not being substantially used in the hybrid decoder. However, the online decode velocity component from LMP pointed less directly towards the target than the spikes’ velocity component (figure 7c). Note that one should not interpret monkey J’s smaller absolute LMP angular error to mean that his LMP signal was more accurate than monkey R‘s; rather, monkey J’s more direct hybrid-driven cursor trajectories (even if they were straighter largely due to better spikes decoding) also brought down the LMP’s absolute errors. These online results contrast with their offline decode counterparts. Offline, decoded LMP aimed the cursor more accurately towards the target than decoded spikes and, not surprisingly, was weighted heavily by the decoder (figure 7(e,f)).

This observation that the LMP aimed the decoded velocity more accurately than spikes towards the target during hand reaches, which are direct and precise, but was less accurate than spikes during closed-loop cursor control, which is noisier and requires more online correction, led us to suspect that the LMP signal was slower than spikes to reflect corrective changes in the animal’s movement intention. To look for evidence of this, we compared the “control lag” for spikes and LMP during closed-loop BMI control by finding the temporal lag that minimized the error angle between each signal’s instantaneous neural push and a vector pointing directly towards the target from the lagged cursor position (figure 7d). In other words, we asked how delayed the response of each signal appeared to be under the (simplistic) assumption that the monkey’s control strategy was always to push the cursor directly towards the target. For both monkeys, the LMP control lag estimated in this manner was twice as long as that of spikes (100 ms versus 50 ms). These error-minimizing lags were the same when closed loop BMI data driven by LMP-only or spikes-only decoders were analyzed (not shown).

**Figure 7.**
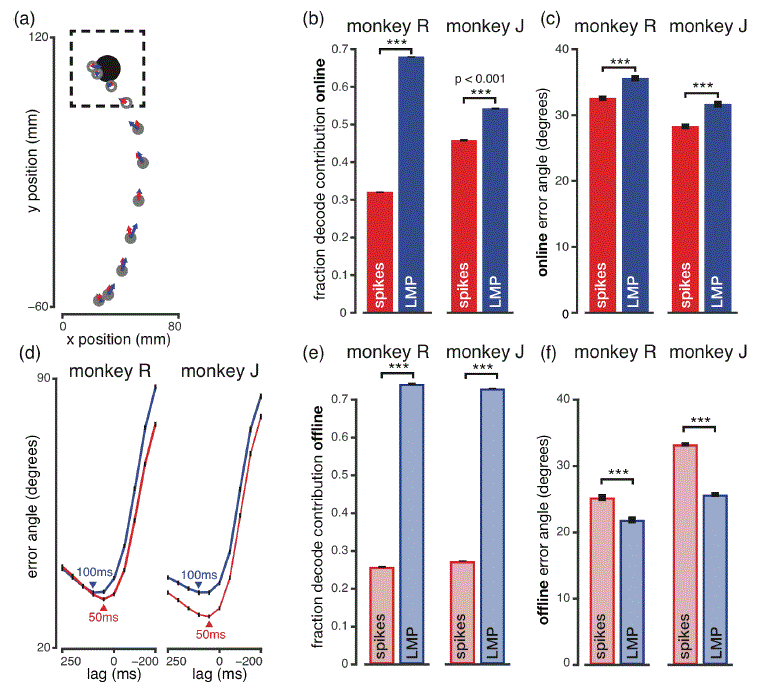
Contribution of spikes and LMP during online and offline hybrid decoding. (a) Example closed-loop BMI trial showing cursor position (grey circles) in successive 50 ms steps as it moved towards the target (black disk). The cursor and target are drawn at ½ size. Instantaneous contributions of the spikes (red) and LMP (blue) signals to decoded velocity are shown as vectors originating from the cursor. Filled cursor positions denote the analysis epoch from 200 ms after target onset until when the cursor first entered the target acceptance region (dashed box). From dataset R.2013.09.27. (b) Mean and SEM of each signal’s relative contribution to the decoded velocity. (c) Angular error was computed between the velocity decoded from each signal and the direction pointing directly towards the target. (d) We repeated the angular error analysis but varied the lag between instantaneous velocity contribution and ‘correct’ angle to estimate control loop lag for each signal. For example, +100 ms lag means that angular error was computed against a vector pointing towards the target from the cursor’s position 100 ms prior. Mean and SEM angular error for the velocity decoded from spikes (red) and LMP (blue) are shown for different lags. (e) Relative decode contribution of spikes and LMP signals in the hybrid decoder was computed as in (b) but using data recorded while the monkey performed the same task with his hand. (f) Error angle was computed as in (c) but using hand control neural data. (b-d) are averaged across all trials from datasets in which the full-channel hybrid decoder was used. (e,f) are averaged across all trials in which the monkeys did the task using hand control.

### Hybrid decoding can rescue performance when channels containing spikes information are removed

In the previous section, we evaluated whether combining LMP with spikes improved BMI performance when both signals were available from all channels. We next investigated whether hybrid decoding could increase the BMI’s robustness to a loss of available spikes information, as would be encountered with degrading arrays. To simulate this scenario, we performed channel-dropping experiments in which we trained hybrid and spikes-only decoders after removing the most contributing spikes channels (see Methods for a description of our channel ranking metric). Closed-loop performance using these channel-dropped decoders was then evaluated on the Continuous Random Target Task (figure 8). Due to the previously described differences in spikes-only, LMP-only, and full-channel hybrid performance between the two monkeys, we treated LMP differently for each monkey’s channel-dropped hybrid decoders. For monkey J, hybrid decoders used spikes from the reduced subset of channels and LMP from all of the channels. This simulates a situation where spikes can no longer be detected on some electrodes, but LMP is still available. For monkey R, we ran a more aggressive channel dropping protocol where both LMP and spikes were removed from dropped channels; this simulates complete failure of the electrodes.

Our motivation for using these two different protocols is as follows. Recall that monkey J’s spikes decoder was much better than his LMP decoder, and his full channel hybrid decoder did not improve upon spikes-only decoding. Monkey J’s arrays recorded good spikes signals from almost every electrode, which resulted in a shallow spikes channel contribution curve (supplementary figure 5). Even with many channels removed, his spikes decoder still outperformed the full-channel LMP decoder (figure 8). Given this, and our earlier observation that adding LMP to much better spikes could be deleterious during closed-loop hybrid decoding, it was not surprising that in pilot studies we found that removing channels for both signals – even when ordered by spikes contribution – still resulted in monkey J’s spikes-only decoders always beating hybrid decoders (up to the point where both failed). We therefore concluded that a more informative monkey J channel dropping experiment would ask: will hybrid decoding ever be helpful if his spikes recording degrades but his mediocre full-channel LMP is still available?

**Figure 8.**
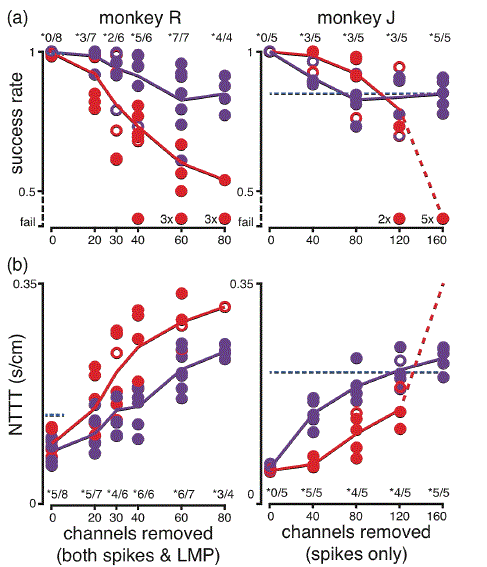
Performance of spikes-only and hybrid decoders after channel dropping. Spikes-only (red) and hybrid spikes + LMP (purple) decoders were trained and evaluated with the specified number of channels removed in a “worst case scenario” by descending order of contribution to the spikes decoder. For monkey R, we removed both spikes and LMP features from these channels. For monkey J only spikes were removed. Spikes and hybrid decoder performances with a given number of channels removed were compared in the same session. Their mean performances are shown as a red and purple point on the plot; filled points represent a significant difference (p < 0.01) for the paired comparison between decoders on that day. Starred fractions summarize how often this difference was significant out of the total number of days when that condition was tested. Solid lines show the grand mean performance of each decoder as a function of channel-drop condition. Days when a decoder ‘failed’ (success rate below 0.5) were not included in this grand mean. Dashed lines extend mean lines into the all-failure conditions for illustrative purposes. Full-channel LMP performance metrics (mean from Figure 6) are shown with horizontal blue lines spanning those conditions where all LMP channels were available to the hybrid decoder. (a) Success rate. (b) NorMalized time to target.

For monkey R, removing only spikes channels (i.e., the monkey J protocol) would not have been very informative given the following observations. Monkey R’s spikes performance was roughly 1.5 times better than his LMP-only performance (a much smaller difference than in monkey J) and relied heavily on a small subset of channels (supplementary figure 5). Once the best few channels were removed, monkey R’s spikes-only decoder became worse than the full-channel LMP decoder (figure 8). Furthermore, from the full-channel hybrid experiments we knew that the two signals already combined beneficially in monkey R’s hybrid decoder, and that this decoder was dominated by LMP even with all spikes channels available. Thus, applying the protocol used in monkey J to monkey R would have rapidly converged to comparing a reduced-channel spikes decoder to a full-channel LMP decoder (plus a minor contribution from remaining spikes). This comparison can be approximated with just a spikes-only channel dropping experiment (which our data does include). By instead comparing reduced channel spikes-only decoders to hybrid decoders in which both signals are removed from dropped channels, we were able to additionally test how much hybrid decoding could rescue performance in the more dire scenario of completely losing electrodes that had the best spikes signals.

We tested monkey J with spikes removed from 0, 40, 80, 120, and 160 channels and LMP still available from all channels. We found that with 40 or 80 channels removed, the hybrid decoder performed worse than a spikes-only decoder (figure 8). Hybrid decoding started to outperform spikes-only decoding with 120 channels removed, performing superiorly in 3/5 sessions. This 120 channels removed condition also corresponds to when spikes-only decoding became worse than the average full-channel LMP-only performance. Once 160 channels were removed, the spikes decoder consistently failed but the hybrid decoder was consistently usable (77-91% trials successful). However, we note that in the conditions where hybrid decoding outperformed spikes-only decoding, it was no better, on average, than LMP-only decoding. Thus, in monkey J hybrid decoding improved robustness to the loss of many spikes channels, but not beyond the improvement afforded by just decoding LMP.

In monkey R, hybrid decoding substantially improved robustness to a complete loss of signals from electrodes containing the best spikes. We compared hybrid and spikes-only decoders with 0, 20, 30, 40, 60, or 80 channels disabled for both spikes and LMP. Hybrid decoders increasingly outperformed their spikes-only counterparts as more channels were removed. This difference was consistently significant (6/6 sessions) once 40 channels were removed; for this condition, success rate was 25% higher and times to target were 38% faster (6 sessions’ grand mean) using the hybrid decoder, not including one session where the spikes decoder failed outright. The hybrid decoder remained consistently usable even with 80 channels removed (77-91% trials successful), whereas the spikes-only decoder started to become unusable once 40 channels were removed. Supplementary movie 2 shows a side-by-side comparison of BMI control during the same session using either a hybrid or spikes-only decoder with 60 channels removed. Three factors contributed to why the improvement afforded by hybrid decoding increased as we removed more channels. First, we dropped channels in order of descending spikes decoder contribution; second, there was a low correlation between how important a given channel was for spikes and LMP decoding (supplementary figure 5 insets); and third, decoder weight was much more evenly distributed across the LMP channels than spikes channels (supplementary figure 5).

## Discussion

The goal of this study was to address the need for motor neural prostheses to be robust to the loss of recordable spikes. The present work substantially advances this goal by demonstrating the effective use of LFP as an alternative or complementary BMI control signal. We decoded a velocity command either solely from multichannel LMP, or from a hybrid control signal consisting of both binned spike counts and LMP. We demonstrated that LMP alone is a viable alternative control signal that enabled effective cursor control in two monkeys. The LMP BMI did not require daily decoder recalibration, and its performance substantially exceeded that of previously reported LFP-driven systems. This is the first study to evaluate BMI control using a single type of LFP feature, in this case the time-varying LFP amplitude – LMP – which is dominated by power below 5 Hz and requires minimal electrical and computational power to extract. We presented new evidence that this signal is not a movement-related artefact by showing that hand velocity could not be decoded from the LMP of sedated monkeys whose arms were passively moved. Finally, we demonstrated for the first time continuous cursor control using a hybrid spikes + LMP decoder. We found that while hybrid decoding did not improve BMI performance when many channels were able to detect spikes, if this spike-based information was removed from some electrodes (simulating sensor degradation), then providing LMP signals to the decoder could rescue performance. Altogether, these results demonstrate that even if multielectrode arrays eventually lose their ability to record spikes on some or even all electrodes, an intracortical BMI may be able to continue to function effectively as long as LMP can still be recorded. The methods described here may therefore extend the useful lifespan of BMI systems, thereby increasing clinical viability.

### Choice of LMP feature for closed-loop BMI use based on offline decode performance

There are myriad ways to process LFP signals into decodable features, and it would not have been experimentally practical to develop and test online BMIs driven by each candidate feature set. Existing offline studies examining the relationship between LFP and reaching kinematics give differing recommendations as to which features are best for BMI use. One study [34] has compared closed-loop performance using LFP power in different frequency bands, but that study reported that the best frequency band was different for each monkey and did not evaluate LMP. Therefore, to maximize our chance of choosing the right LFP feature(s), we first performed our own offline comparisons which found that LMP was the best of the tested LFP-derived features. This is in accordance with past studies showing that LMP recorded from primary motor and premotor cortices is informative about kinematics [17,18,20-22,24,58,59]. Interestingly, both our study and previous studies [19,25-27] found that power in low-frequency LFP bands is a poor decode feature. This suggests that even though low-frequency components dominate the LMP, the loss of phase information when estimating low-frequency power results in a loss of useful information.

The second best LFP feature we tested, power in the 50-300 Hz band, performed considerably worse than LMP in our offline decode comparison. This contrasts with [21,22] which found that high-frequency LFP power was comparable to or better than LMP. Such differences might be expected if the relative information contained in LMP versus high frequency LFP power is sensitive to precisely where the signals are recorded. We are unaware of any study that methodically mapped how the quality of LMP information changes as a function of recording location, but one study [27] showed that decoding performance using higher frequency spectral features depends strongly on cortical recording depth. Similar studies to optimize recording location for LMP-based decoding would be very useful.

We were surprised to find that our offline decode accuracy using LMP was comparable to decoding multiunit spikes. Although previous studies have shown better or equal offline decoding using LFP features rather than spikes from a small number of channels [18,21,23-25,59], many of the same studies [21,23,24,59], and others [22,58], reported that at higher channel counts, multichannel spike activity is more informative than multichannel LFP. Our 192-channel study falls within the upper channel count range of these past reports. However, differences in electrode placement, signal processing and decoding techniques, and the nature of the kinematics being decoded obfuscate direct comparisons between studies and may explain why our offline decode of velocity was better using LFP than using spikes. Consistent with previous reports [18,23,58,59], we found that offline decoding of both LFP and spikes together led to a modest but significant accuracy improvement. This improvement was greater using LMP than 50-300 Hz power. Although our results do not speak to this directly, one reason why high frequency LFP power may not add much information to what can already be decoded from multiunit spikes is that this signal may itself be contaminated by spiking activity [28,60].

Taken together, our offline decode results led us to conclude that LMP was the most promising candidate LFP feature both as an alternative and complementary signal to spikes for BMI control. These data also address several concerns that have been raised about the utility of low-frequency LFP signals for BMI use. One such concern is that decoding low-frequency LFP may be hindered by high correlations of both signal and noise between electrodes [17,22,23,59]. While we did observe that LMP from nearby electrodes had similar tuning, there were a variety of PDs across the two 96-electrode arrays and we were able to accurately decode single-trial velocity. A second concern is that the signal is an artefact caused by the animal’s movement. Our sedated animal decode control experiment adds a new piece of evidence on top of the already-persuasive case made by prior studies [32,59] that low-frequency LFP is not merely a mechanical or electrical artefact. A third concern is that LMP would be ill-suited to serve as a neural prosthetic signal if it lags behind kinematics (i.e., contains mainly sensory information); this has been reported in motor [59] and parietal [28] cortices. Contrary to these studies, we found that kinematics were best decoded when LMP was time-shifted to precede kinematics by approximately 100 ms. Our results may differ from [28] because they recorded from a different cortical region, and from [59] because they appear to have have recorded from an area with more sensory information (they also found that most single units’ optimal decoding lags followed kinematics). Our results are consistent with a different set of studies which found that that LMP is tuned before movement onset [18,24,32].

### Closed-loop LMP-driven BMI performance

This study demonstrates that decoding LMP from two multielectrode arrays can enable effective BMI cursor control. It adds to the small existing body of closed-loop LFP-driven cursor control work [33,34] by demonstrating improved performance while also characterizing the quality of control using a single LFP-derived feature (LMP). Differences in monkeys and signal recording quality inevitably limit how much can be inferred by comparing the performance of different BMI methods across studies. Nonetheless, we made a best effort to directly compare the performance of our LMP-driven BMI to that of two previous closed-loop LFP-driven BMI studies on closely matched tasks. We observed target acquisition rates roughly 50% higher than [34] and over twice as fast as [33]. Thus, the work described here represents, to the best of our knowledge, the highest-performing LFP-driven BMI system to date. We believe that two design features contributed to better performance using our BMI system. First, we decoded neural activity from more electrodes than previous studies. We recorded from two 96-electrode arrays (one in M1 and one in PMd), whereas So and colleagues [34] decoded LFP from twenty electrodes randomly chosen from a 128-electrode array spanning M1 and PMd, and Flint and colleagues [33] recorded from a single 96-channel array in M1. Second, we decoded LMP rather than spectral power, which according to our offline decoding analysis better predicts reach velocity. Flint and colleagues also decoded LMP as one of their features, but did not implement something akin to our half-wave rectification. If either of their monkeys exhibited the same deleterious LMP after-potential that we observed in our monkey R (perhaps partially compensated for by other decoded LFP power features), this may have limited performance. So and colleagues’ decoders used multiple LFP power bands between 0 and 150 Hz, but did not decode LMP.

This substantially better LFP-driven performance serves as an important existence proof that high-quality BMI control is possible using alternative (non-spike) signals from chronically implanted multielectrode arrays. Importantly, however, we found that our LMP-driven BMI performance was not as good as spike-driven performance using the state-of-the-art FIT-KF algorithm [49]. Thus, while our results demonstrate that LMP is a viable alternative control signal to spikes, we do not suggest employing LFP-only decoding if good spikes are available. We do note, however, that the low sampling rate and minimal signal processing required to extract LMP, combined with its reasonable performance as a closed-loop control signal, may make LMP attractive for power-or bandwidth-constrained applications such as fully implanted BMI systems (these constraints are overviewed in e.g., [61]). The low-pass filtering properties used to extract LMP in this study can be approximated with a passive analog filter followed by half-wave rectification via a very low power analog rectification circuit. Downstream processing would only need to sample the resulting voltage 40 times per second (the decode update time used in this study). This contrasts dramatically with the order 103 samples per second used to identify single unit spikes. Generally speaking, recording sorted spikes has the highest power demands, followed by threshold crossings and higher frequency LFP band power, and finally LMP.

A desirable property of a neural prosthetic control signal is not just the performance that it affords, but also its stability. BMI use without daily decoder recalibration has been shown when decoding well-isolated single units [2], multiunit spiking [16,33], and a broad set of LFP-derived features including both LMP and higher frequency power [33]. We found that our LMP-only decoder can also be used without retraining for at least three weeks. Unlike in [33], we did not see long-term performance improvement when using a fixed decoder. This may be because we did not hold the decoder stable for as many days or for as long during each day, or because the higher performance in our study reduced the pressure on the animal to improve his performance.

Our study was similar to that of Flint and colleagues [33], but differed from the So and colleagues study [34], in that we allowed the monkeys to continue to move their arm during BMI use. This arm-free BMI animal model is better suited for evaluating biomimetic decoders trained using arm reaching data [44], and unsurprisingly we observed that monkeys continued to move their arms when using the LMP-driven BMI. In this regard LMP-driven BMI use is similar to spike-driven control (both in this and past studies such as [3]), during which monkeys typically continue to move their arms if allowed to. As with any neural prosthetics method tested using an able-bodied animal model, it remains to be seen how well our approach will work in paralyzed or amputee patients. While we are encouraged by a recent report of movement intention-evoked transient low-frequency LFP modulation in humans with tetraplegia implanted with similar arrays [30], validation of our LMP decoding approach will require closed-loop evaluation of this method in clinical trial participants.

### Hybrid decoding using both LMP and spikes

Decoding spikes or LFP need not be an either-or decision. While it has often been predicted that hybrid decoding of both signals could improve BMI performance, this is the first closed-loop study to decode continuous kinematics from spikes and LFP together. In doing so, we faced two decisions: how should we combine these two signals in a single decoder, and what spikes-only decoder should we compare the resulting hybrid decoder against? We felt that it would be most impactful to try to improve performance and robustness above that of a current state-of-the-art spikes decoder. We therefore used as our baseline the FIT-KF algorithm [49] and then extended it to decode both spikes and LMP. When both spikes and LMP were decoded from all available electrodes, hybrid decoding enabled a modest improvement in closed-loop performance in monkey R, but was marginally worse than spikes-only decoding in monkey J. An analysis of each signal’s contribution to the closed-loop velocity decode provides insight into why adding LMP did not improve closed-loop performance as much as might have been expected from offline analyses: when the hybrid decoder is fit, LMP features are heavily weighted because collectively they are as well as or better correlated than spikes with the open-loop training data kinematics. During closed-loop BMI use, however, even modest decode errors must be corrected in order to acquire the target; consequently, the longer control lag characteristic of LMP leads to worse control. As a result, the LMP’s contribution to a hybrid decoder with excellent spikes signals does little to improve – and can even worsen – closed-loop cursor control. Consequently, achieving a benefit from online hybrid decoding is not a given, but rather will depend on the relative *closed-loop* quality of the spikes-and LMP-signals.

We note two caveats related to our design choices of building the hybrid decoder by extending the FIT-KF spikes decoder and then comparing it against the spikes-only variant. First, while combining the spikes and LMP Kalman filters was a principled application of existing high-performing decoding techniques to the hybrid decoding problem, we make no claim that the resulting decoder – which determines the relative weighting of spikes and LMP features based on how well each signal regressed against training data kinematics and how the resulting residuals covaried – is optimal for closed-loop BMI use. We view this work as an encouraging first foray into hybrid decoding but recognize a need for future work looking at whether there are better ways to combine spikes and LFP signals. For example, there may be an opportunity to improve performance by taking into account the different control loop latencies of LMP compared to spikes, or by decreasing the decoder weights assigned to LMP features to account for their inferior closed-loop utility. Second, by comparing the hybrid decoder to what was already a very high performing spikes-only decoder, we set a high bar for performance improvement. Monkey J, in particular, had many good channels with spikes and controlled the cursor almost as well with the full-channel FIT-KF spikes decoder as with his hand. With these caveats in mind, the present results suggest that when many channels with spikes information are available, hybrid decoding offers little to no improvement over state-of-the-art spikes-only decoding.

### Decoding LMP to mitigate losing spikes channels

Thus far we have discussed how LMP and hybrid decoding compares to spikes-based decoding in the case of arrays with excellent (monkey J) or mediocre (monkey R) spike signals. But what about when spike signal quality is poor and insufficient for accurate BMI control? This is oftentimes the case when arrays degrade over the years following implantation [14,29] and is one of the major impediments to long-lasting, clinically viable neural prostheses. We experimentally simulated this situation with channel dropping experiments, and found that the performance drop resulting from losing spikes information can be mitigated by making use of LMP. The degree to which LMP provides robustness to spikes loss depends on the relative quality of the LMP and remaining spikes, as we will now describe.

In Monkey J, LMP was in effect a BMI control signal of last resort. This subject had excellent spike signals on most channels, which enabled much better cursor control using a spikes-driven BMI than with an LMP-driven BMI. Only after a majority of spikes channels were dropped did his spikes-only control become worse than full-channel LMP-only control. At that point, both hybrid decoding and LMP-only decoding (both of which had LMP available from all channels) provided usable cursor control, whereas the spikes-only decoder failed. Importantly, across the channel-dropping conditions we tested, the hybrid decoder did not outperform whichever individual signal type (spikes-only or LMP-only) worked best. It is somewhat counterintuitive that adding additional LMP information to monkey J’s channel-dropped spikes signal actually reduced performance in the conditions where spikes still outperformed full-channel LMP. This (suboptimal) hybrid decoding result can be understood in light of the previously discussed observation that LMP and spikes were comparable for offline decoding but that LMP provided worse closed-loop control. When monkey J’s most contributing spikes channels were removed, the regression to fit a hybrid decoder increasingly weighed the LMP over the remaining spikes because of LMP’s high offline explanatory power. This resulted in a hybrid decoder that was (overly) dominated by the inferior LMP signal rather than the more controllable remaining spikes.

In contrast, utilizing LMP improved monkey R’s BMI control even when spikes were available from all channels, and the LMP became increasingly important for maintaining performance as spike signals were lost. Monkey R had worse spikes-driven control and better LMP-driven control than monkey J, such that full-channel LMP decoding outperformed spikes-only decoding once twenty or more spikes channels were lost. This suggests that when spikes’ performance advantage over LMP is not large (and is tenuous insofar as it depends on a handful of key channels), a modest degree of array spikes signal decay will result in LMP decoding becoming a preferred alternative to spikes decoding. Furthermore, monkey R’s hybrid decoder outperformed the spikes-only decoder across all channel drop conditions tested, demonstrating that when closed-loop LMP-driven control is comparable to (or better than) spikes-driven control, LMP can complement spikes and improve BMI performance.

Taken together these results provide experimental validation that utilizing LFP (in this study, specifically the LMP feature) as a BMI control signal can increase a neural prosthesis’ robustness to the loss of spikes signal. At a minimum,

LMP can provide an alternative control signal to spikes (as in monkey J’s case), which is useful because there is considerable experimental evidence suggesting that arrays’ LFP signals persist longer than spikes [22, 23, 28, 30, 39, 40]. In the more fortuitous case where LMP-driven performance is quite good (such as in monkey R), LMP can be used not just as an alternative to spikes, but as a complementary signal for a hybrid decoder that both improves maximal performance, and slows down the rate at which performance decays when spikes signals are lost. We are particularly encouraged that in this study, decoding LMP enabled effective cursor control in both monkeys under conditions where spikes-only decoding was non-functional.

There are several channel-dropping experiment permutations that this study did not explore but which are worth considering. We did not test monkey R in the case that spikes were lost from dropped channels but LMP was still available from all channels. However, the observation that his full-channel LMP-only decoder was better than any of the channel-dropped spikes-only decoders, and that LMP and spikes combined beneficially in the full-channel hybrid case, strongly predicts that hybrid decoding would be even better if more LMP channels are available. Since the main goal of our study was to use available LMP signals to increase robustness to a potential loss of spikes signals, we also did not explicitly explore how these decoders would perform when, instead, LMP channels are lost (i.e., LMP-only channel-dropping experiments, or hybrid-decoder channel dropping ordered by LMP contribution). Based on the high correlations between LMP on nearby channels and the shallow LMP decoder contribution curves, we predict that such experiments would find that LMP and hybrid decoders would degrade slower (as a function of number of channels removed) than spikes-driven decoders. This prediction remains to be tested in future closed-loop experiments.

### Differences between offline and closed-loop results

Our experiences carrying out this work add to the growing body of literature reporting that offline decoding results do not always predict closed-loop BMI performance (e.g., [46,62]). Given the field’s heavy reliance on offline decode analyses, we believe it is worthwhile to enumerate these differences in order to draw attention to factors that may affect how offline performance translates to closed-loop results. Although our offline metrics showed that decoding LMP was comparable to or even better than decoding spikes, during closed-loop evaluation spikes-driven cursor control was consistently superior. Qualitatively, the LMP-only decoder appeared more ‘ballistic’ and less precisely controllable than the spikes-only decoder: if the monkey missed the target with his initial reach, his corrective response was slower and less smooth. This impression is supported by several observations: LMP-only decoding had longer times to target and dial-in time; during hybrid decoding, the decoded LMP aimed the cursor less accurately than spikes did; and the estimated control loop lag was longer for LMP than for spikes. We therefore attribute LMP’s reduced closed-loop controllability to this signal being slower to reflect changing movement intention – a limitation that is likely to be shared by other low-frequency signals. To better predict such offline versus closed-loop differences in the future, we suggest that the datasets used to evaluate candidate BMI control signals include conditions in which rapid reach corrections are made. This would enable estimation of the candidate signal’s control lag. LMP’s inferior closed-loop controllability also explains a second difference between offline and closed-loop decoding: offline, hybrid spikes + LMP decoding helped in both monkeys, but during closed-loop control this benefit was only realized in monkey R.

A third example is that decoding LMP without half-wave rectification worked well offline, but in one monkey halfwave rectification was needed to enable closed-loop performance. This signal processing step removed an LMP ‘afterpotential’ (visible at t = 0.6 s in figure 1d) which occurred shortly after the monkey reached towards the target. Without rectification, this afterpotential generated a substantial cursor velocity during the time period when the monkey was typically trying to keep the cursor still over the target. After we discovered this problem during closed-loop BMI testing, we looked more closely at our offline decode results and noticed that the decoded velocities did slightly increase at the end of the target hold period. However, this deleterious effect was poorly captured in our offline decode metrics because the afterpotential happened near the end of the trial. Consequently, the decoded hand velocity was just beginning to increase when the trial ended, preventing further accumulation of decode error. The next trial’s decode then began with the Kalman state reset to zero (a common practice to prevent offline decoding errors from accumulating over the course of many trials), minimizing the afterpotential’s effect on this next trial. In short, the trial structure of the data we used for the offline study largely hid the problem, but in closed-loop the afterpotential’s effect was strong enough to dramatically hinder target acquisition.

Offline studies are invaluable for suggesting promising BMI design approaches, but these differences underscore the need for careful closed-loop validation. Thus, while we specifically investigated closed-loop LMP-driven performance because this LFP feature worked best offline, our results do not rule out that other features cannot yield even better closed-loop performance. The nascent subfield of LFP-driven BMIs would benefit from more studies evaluating other choices of LFP features and decoding algorithms in closed-loop experiments.

## Acknowledgements

This work was supported by National Science Foundation’s Graduate Research Fellowships (SDS, JCK) and IGERT award 0734683 (SDS), Stanford Medical Scientist Training Program (PN), NIH Pioneer Award 8DP1HD075623 (KVS), NIH T-RO1 award NS076460 (KVS), and DARPA REPAIR award N66001-10-C-2010 (KVS).

We thank M. Mazariegos, J. Aguayo, M. Wechsler, C. Sherman, E. Morgan, & L. Yates for expert surgical assistance & veterinary care, C. Chandrasekaran for Simulink real-time code to access LFP data, B. Oskotsky for IT support, and B. Davis, & E. Casteneda for administrative assistance.

## References

[1] Serruya M, Hatsopoulos N G, Paninski L, Fellows M R and Donoghue J P 2002 Instant neural control of a movement signal Nature 416 141–2

[2] Ganguly K and Carmena J M 2009 Emergence of a stable cortical map for neuroprosthetic control PLoS Biol. 7 e1000153

[3] Gilja V, Nuyujukian P, Chestek C A, Cunningham J P, Yu B M, Fan J M, Churchland M M, Kaufman M T, Kao J C, Ryu S I and Shenoy K V 2012 A high-performance neural prosthesis enabled by control algorithm design Nat. Neurosci. 15 1752–7

[4] Carmena J M, Lebedev M A, Crist R E, O‘Doherty J E, Santucci D M, Dimitrov D F, Patil P G, Henriquez C S and Nicolelis M A L 2003 Learning to control a brain-machine interface for reaching and grasping by primates PLoS Biol. 1 193–208

[5] Velliste M, Perel S, Spalding M C, Whitford A S and Schwartz A B 2008 Cortical control of a prosthetic arm for self-feeding Nature 453 1098–101

[6] Ethier C, Oby E, Bauman M and Miller L E 2012 Restoration of grasp following paralysis through brain-controlled stimulation of muscles Nature 1–4

[7] Bacher D, Jarosiewicz B, Masse N Y, Stavisky S D, Simeral J D, Newell K, Oakley E M, Cash S S, Friehs G and Hochberg L R 2014 Neural Point-and-Click Communication by a Person With Incomplete Locked-In Syndrome Neurorehabil. Neural Repair

[8] Hochberg L R, Bacher D, Jarosiewicz B, Masse N Y, Simeral J D, Vogel J, Haddadin S, Liu J, Cash S S, vander Smagt P and Donoghue J P 2012 Reach and grasp by people with tetraplegia using a neurally controlled robotic arm Nature 485 372–5

[9] Collinger J L, Wodlinger B, Downey J E, Wang W, Tyler-Kabara E C, Weber D J, McMorland A J C, Velliste M, Boninger M L and Schwartz A B 2013 High-performance neuroprosthetic control by an individual with tetraplegia. Lancet 381 557–64

[10] Ryu S I and Shenoy K V 2009 Human cortical prostheses: lost in translation? Neurosurg. Focus 27 E5

[11] Simeral J D, Kim S-P, Black M J, Donoghue J P and Hochberg L R 2011 Neural control of cursor trajectory and click by a human with tetraplegia 1000 days after implant of an intracortical microelectrode array J. Neural Eng. 8 025027

[12] Krüger J, Caruana F, Volta R D and Rizzolatti G 2010 Seven years of recording from monkey cortex with a chronically implanted multiple microelectrode. Front. Neuroeng. 3 6

[13] Chestek C A, Gilja V, Nuyujukian P, Foster J D, Fan J M, Kaufman M T, Churchland M M, Rivera-Alvidrez Z, Cunningham J P, Ryu S I and Shenoy K V 2011 Long-term stability of neural prosthetic control signals from silicon cortical arrays in rhesus macaque motor cortex J. Neural Eng. 8 045005

[14] Barrese J C, Rao N, Paroo K, Triebwasser C, Vargas-Irwin C, Franquemont L and Donoghue J P 2013 Failure mode analysis of silicon-based intracortical microelectrode arrays in non-human primates. J. Neural Eng. 10 066014

[15] Fraser G W, Chase S M, Whitford A and Schwartz A B 2009 Control of a brain-computer interface without spike sorting. J. Neural Eng. 6 055004

[16] Nuyujukian P, Kao J C, Fan J M, Stavisky S D, Ryu S I and Shenoy K V 2014 Performance sustaining intracortical neural prostheses. J. Neural Eng. 11 066003

[17] O’Leary J G and Hatsopoulos N G 2006 Early Visuomotor Representations Revealed From Evoked Local Field Potentials in Motor and Premotor Cortical Areas J. Neurophysiol. 96 1492–506

[18] Mehring C, Rickert J, Vaadia E, Cardoso de Oliveira S, Aertsen A and Rotter S 2003 Inference of hand movements from local field potentials in monkey motor cortex Nat. Neurosci. 6 1253–4

[19] Scherberger H, Jarvis M R and Andersen R A 2005 Cortical local field potential encodes movement intentions in the posterior parietal cortex. Neuron 46 347–54

[20] Rickert J, Oliveira S, Vaadia E, Aersten A, Rotter S and Mehring C 2005 Encoding of Movement Direction in Different Frequency Ranges of Motor Cortical Local Field Potentials J. Neurosci. 25 8815–24

[21] Bansal A K, Truccolo W, Vargas-Irwin C E and Donoghue J P 2011 Decoding 3D reach and grasp from hybrid signals in motor and premotor cortices: spikes, multiunit activity, and local field potentials J. Neurophysiol. 107 1337–55

[22] Flint R D, Lindberg E W, Jordan L R, Miller L E and Slutzky M W 2012 Accurate decoding of reaching movements from field potentials in the absence of spikes J. Neural Eng. 9 046006

[23] Hwang E J and Andersen R A 2013 The utility of multichannel local field potentials for brain-machine interfaces. J. Neural Eng. 10 046005

[24] Flint R D, Ethier C, Oby E R, Miller L E and Slutzky M W 2012 Local field potentials allow accurate decoding of muscle activity. J. Neurophysiol. 4653

[25] Pesaran B, Pezaris J S, Sahani M, Mitra P P and Andersen R A 2002 Temporal structure in neuronal activity during working memory in macaque parietal cortex Nat. Neurosci. 5 805–11

[26] Zhuang J, Truccolo W, Vargas-Irwin C and Donoghue J P 2010 Decoding 3-D reach and grasp kinematics from high-frequency local field potentials in primate primary motor cortex. ?IEEE Trans. Biomed. Eng. 57 1774–84

[27] Markowitz D A, Wong Y T, Gray C M and Pesaran B 2011 Optimizing the Decoding of Movement Goals from Local Field Potentials in Macaque Cortex J. Neurosci. 31 18412–22

[28] Asher I, Stark E, Abeles M and Prut Y 2007 Comparison of Direction and Object Selectivity of Local Field Potentials and Single Units in Macaque Posterior Parietal Cortex During Prehension J. Neurophysiol. 3684–95

[29] Wang D, Zhang Q, Li Y, Wang Y, Zhu J, Zhang S and Zheng X 2014 Long-term decoding stability of local field potentials from silicon arrays in primate motor cortex during a 2D center out task. J. Neural Eng. 11 036009

[30] Perge J A, Zhang S, Malik W Q, Homer M L, Cash S, Friehs G, Eskandar E N, Donoghue J P and Hochberg L R 2014 Reliability of directional information in unsorted spikes and local field potentials recorded in human motor cortex. J. Neural Eng. 11 046007

[31] Kennedy P R, Kirby M T, Moore M M, King B and Mallory A 2004 Computer Control Using Human Intracortical Local Field Potentials IEEE Trans. Neural Syst. Rehabil. Eng. 12 339–44

[32] Hwang E J and Andersen R A 2009 Brain control of movement execution onset using local field potentials in posterior parietal cortex. J. Neurosci. 29 14363–70

[33] Flint R D, Wright Z A, Scheid M R and Slutzky M W 2013 Long term, stable brain machine interface performance using local field potentials and multiunit spikes J. Neural Eng. 10 056005

[34] So K, Dangi S, Orsborn A L, Gastpar M C and Carmena J M 2014 Subject-specific modulation of local field potential spectral power during brain-machine interface control in primates J. Neural Eng. 11 026002

[35] Belitski A, Gretton A, Magri C, Murayama Y, Montemurro M A, Logothetis N K and Panzeri S 2008 Low-frequency local field potentials and spikes in primary visual cortex convey independent visual information J. Neurosci. 28 5696–709

[36] Katzner S, Nauhaus I, Benucci A, Bonin V, Ringach D L and Carandini M 2009 Local origin of field potentials in visual cortex. Neuron 61 35–41

[37] Buzsáki G, Anastassiou C A and Koch C 2012 The origin of extracellular fields and currents – EEG, ECoG, LFP and spikes Nat. Rev. Neurosci. 13 407–20

[38] Einevoll G T, Kayser C, Logothetis N K and Panzeri S 2013 Modelling and analysis of local field potentials for studying the function of cortical circuits Nat. Rev. Neurosci. 14 770–85

[39] Heldman D A, Wang W, Chan S S and Moran D W 2006 Local Field Potential Spectral Tuning in Motor Cortex During Reaching IEEE Trans. Neural Syst. Rehabil. Eng. 14 180–3

[40] Bokil H S, Pesaran B, Andersen R A and Mitra P P 2006 A method for detection and classification of events in neural activity IEEE Trans. Biomed. Eng. 53 1678–87

[41] Stavisky S D, Kao J C, Nuyujukian P, Ryu S I and Shenoy K V 2014 Hybrid Decoding of Both Spikes and Low-Frequency Local Field Potentials for Brain-Machine Interfaces 36th Annual International Conference of the IEEE Engineering in Medicine and Biology Society (Chicago, IL) pp 3041–4

[42] Santhanam G, Ryu S I, Yu B M, Afshar A and Shenoy K V 2006 A high-performance brain-computer interface Nature 442 195–8

[43] Shenoy K V. and Carmena J M 2014 Combining Decoder Design and Neural Adaptation in Brain-Machine Interfaces Neuron 84 665–80

[44] Nuyujukian P, Fan J M, Gilja V, Kalanithi P S, Chestek C A and Shenoy K V 2011 Monkey Models for Brain-Machine Interfaces: The Need for Maintaining Diversity 33rd Annual International Conference of the IEEE Engineering in Medicine and Biology Society (Boston, MA)

[45] Dangi S, So K, Orsborn A L, Gastpar M C and Carmena J M 2013 Brain-machine interface control using broadband spectral power from local field potentials 35th Annual International Conference of the IEEE EMBS (Osaka, Japan) pp 285–8

[46] Cunningham J P, Nuyujukian P, Gilja V, Chestek C A, Ryu S I and Shenoy K V 2010 A Closed-Loop Human Simulator for Investigating the Role of Feedback-Control in Brain-Machine Interfaces. J. Neurophysiol.

[47] Wu W, Gao Y, Bienenstock E, Donoghue J P and Black M J 2006 Bayesian Population Decoding of Motor Cortical Activity Using a Kalman Filter Neural Comput. 118 80–118

[48] Kim S-P, Simeral J D, Hochberg L R, Black M J, General M and Systems C N 2008 Neural control of computer cursor velocity by decoding motor cortical spiking activity in humans with tetraplegia J. Neural Eng. 5 455–76

[49] Fan J M, Nuyujukian P, Kao J C, Chestek C A, Ryu S I and Shenoy K V 2014 Intention estimation in brain-machine interfaces J. Neural Eng. 11 016004

[50] Kuo B C 1987 Automatic Control Systems (Englewood Cliffs: Prentice-Hall, Inc.)

[51] MacKenzie I S and Ware C 1993 Lag as a determinant of human performance in interactive systems Proceedings of the SIGCHI conference on Human factors in computing systems – CHI’93 (New York, New York, USA: ACM Press) pp 488–93

[52] Willett F R, Suminski A J, Fagg A H and Hatsopoulos G 2012 Compensating for Delays in Brain-Machine Interfaces by Decoding Intended Future Movement 4087–90

[53] Suminski A J, Tkach D C, Fagg A H and Hatsopoulos N G 2010 Incorporating Feedback from Multiple Sensory Modalities Enhances Brain-Machine Interface Control J. Neurosci. 30 16777–87

[54] Fitts P M 1954 The information capacity of the human motor system in controlling the amplitude of movement. J. Exp. Psychol. 47 381–91

[55] Douglas S, Kirkpatrick A and MacKenzie I 1999 Testing pointing device performance and user assessment with the ISO 9241, Part 9 standard Proc. ACMSIGCHI Conf. Hum. Factors Comput. Syst. 215–22

[56] MacKenzie I S and Buxton W 1992 Extending fitts’ law to two-dimensional tasks Proc. SIGCHI Conf. Hum. Factors Comput. Syst. 219–26

[57] Malik W Q, Truccolo W, Brown E N and Hochberg L R 2011 Efficient decoding with steady-state Kalman filter in neural interface systems IEEE Trans. Neural Syst. Rehabil. Eng. 19 25–34

[58] Stark E and Abeles M 2007 Predicting movement from multiunit activity. J. Neurosci. 27 8387–94

[59] Bansal A K, Vargas-Irwin C E, Truccolo W and Donoghue J P 2011 Relationships among low-frequency local field potentials, spiking activity, and three-dimensional reach and grasp kinematics in primary motor and ventral premotor cortices. J. Neurophysiol. 105 1603–19

[60] Waldert S, Lemon R N and Kraskov A 2013 Influence of spiking activity on cortical local field potentials J. Physiol. 591 5291–303

[61] Dethier J, Nuyujukian P, Ryu S I, Shenoy K V and Boahen K 2013 Design and validation of a real-time spiking-neural-network decoder for brain-machine interfaces J. Neural Eng. 10 036008

[62] Chase S M, Schwartz A B and Kass R E 2009 Bias, optimal linear estimation, and the differences between open-loop simulation and closed-loop performance of spiking-based brain-computer interface algorithms. Neural Networks 22 1203–13

